# Fluorescence *in situ* hybridization (FISH) and detection of Bacteria and Archaea cells in fecal samples

**DOI:** 10.1101/2025.10.02.679970

**Authors:** Maria Nejjari, Michel cloutier, Guylaine Talbot, Martin Lanthier

## Abstract

Oligonucleotide probes have been used to detect bacteria and archaea that colonize the cattle and pig digestive system.

The Fluorescent *in situ* hybridization (FISH) is a molecular biology technique that uses oligonucleotides of 15 to 25 nucleotides of length associated with a fluorescent molecule.

In microbiology, the FISH technique utilizes probes targeting ribosomal RNA (rRNA). It is one amongst others, a staining technique that allows the identification, detection and quantification of microorganisms without prior cultivation by means of epifluorescence and confocal laser scanning microscopy (CLSM).

In this study, we describe the usage of the confocal laser scanning microscopy coupled with fluorescence in situ hybridization (FISH), in order to detect and quantify of bacteria and archaea in fecal samples from cattles’ manure and swine slurry.

## Introduction

Microbial communities in the gastrointestinal tract of ruminants (eg. cattles, sheep) and non-ruminants (eg. Swine slurry, Human) play a crucial role in host health and performance (Mohapatra, 2008). This role appears during the provision of nutrients (Cummings *et al*. 1997 and Topping *et al*. 2001), stimulation of immune system response (Kimura *et al*. 1997), metabolism of toxins (Macfarlane *et al*. 1997 et Salyers *et al*. 1984) and gene expression in host tissue (Tran *et al*. 1998 and Hooper *et al*. 2001).

The most abundant microorganisms in animal intestines are archeae and bacteria (called also eubacteria). Fecal material encloses panoply of microorganisms. Probiotics form a defense barrier for the host. Pathogens like *E*.*coli O157, Listeria, Salmonella* can easily contaminate water streams, agricultural soils and by consequence food products which represent a source of infection to humans and animals.

Methanogens are archeae found in fecal material of animals. Species like *Methanoculleus spp*. and *Methanobacterium spp*., amongst others, are very interesting to study especially that they represent a potential interest in Methane production as a source of renewable energy.

In order to study and quantify the cultivable predominant species in the microbial composition of the gastrointestinal tract, previous researches concentrated on different culture techniques using fecal samples (Bottari *et al*. 2006; Finegold *et al*. 1983; Holdeman *et al*. 1976; Moore *et al*. 1995). However, those techniques are very laborious, time consuming, susceptible to statistical and experimental errors (Moter et Gobel 2000) and need microorganism isolation from its matrix of origin in order to study it (Bottari *et al*. 2006).

During last years, there were great advances in the analysis of complex microbial ecosystems using molecular techniques (Amann *et al*. 1990; Raskin *et al*. 1994; Stahl *et al*. 1988). *In situ* hybridization is used as a molecular technique based on ribosomal RNA (rRNA) (Barc *et al*. 2004).

In microbiology, the FISH technique relies on the use of oligonucleotides with a length of 15 to 25 nucleotides and that are associated with a fluorescent molecule. These oligonucleotides, named “Molecular probes”, are generally complementary to the 16S or 23S rRNA. For the FISH technique, it is important to use 20 nucleotides length probes (Lanthier 2004). The FISH technique helps in recognizing the phylogenic identity, morphology, abundance and spatial arrangement of microorganisms in their original matrix (Amann *et al*. 1995).

The success of FISH experiments is generally measured as the fraction of detectable cells in order to determine the microbial communities. This fraction is a comparison between the numbers of microorganisms detected with archeae or bacteria specific fluorescent probes and the total microorganisms labeled with archeae and bacteria specific probe (Hicks *et al*. 1992).

In this, we describe the usage of confocal laser scanning microscope (CLSM) coupled with fluorescence in situ hybridization (FISH) to detect and quantify bacteria and archeae, using 16S rRNA-targeted oligonucleotides probes, in fecal samples issued from cattles’ manure and swine slurry.

## Materials and Methods

### Source of the analyzed samples

bacterial strains have been provided by the Dairy and Swine Research Development Centre, Agriculture and Agri-food Canada (Lennoxville, Québec, Canada). These bacterial strains have been obtained from fecal samples issued from cattle manure and swine slurry (Table 1). The collect of these samples has been done at 3 different pit levels A, B and C with same height. Two collect depth levels have been taken into consideration, one collection level just under the pit’s surface and the second one at the bottom of the pit.

**Table 1.** Fecal samples issued from cattle manure and swine slurry.

| sample. # | identification | Pit's Source & # | Collection depth | Collection # | Collection place having same level | Date |
| --- | --- | --- | --- | --- | --- | --- |
| 73 | F1AH1-T3 | Swine slurry #1 | Under the pit surface | T3 | A | 19-04-2011 |
| 74 | F1AH3-T3 | Swine slurry #1 | At the pit's bottom | T3 | A | 19-04-2011 |
| 75 | F1BH1-T3 | Swine slurry #1 | Under the pit surface | T3 | B | 19-04-2011 |
| 76 | F1BH3-T3 | Swine slurry #1 | At the pit's bottom | T3 | B | 19-04-2011 |
| 77 | F1CH1-T3 | Swine slurry #1 | Under the pit surface | T3 | C | 19-04-2011 |
| 78 | F1CH3-T3 | Swine slurry #1 | At the pit's bottom | T3 | C | 19-04-2011 |
| 79 | F2AH1-T3 | Swine slurry #2 | Under the pit surface | T3 | A | 19-04-2011 |
| 80 | F2AH3-T3 | Swine slurry #2 | At the pit's bottom | T3 | A | 19-04-2011 |
| 81 | F2BH1-T3 | Swine slurry #2 | Under the pit surface | T3 | B | 19-04-2011 |
| 82 | F2BH3-T3 | Swine slurry #2 | At the pit's bottom | T3 | B | 19-04-2011 |
| 83 | F2CH1-T3 | Swine slurry #2 | Under the pit surface | T3 | C | 19-04-2011 |
| 84 | F2CH3-T3 | Swine slurry #2 | At the pit's bottom | T3 | C | 19-04-2011 |
| 85 | F3AH1-T3 | Cattle pit #1 | Under the pit surface | T3 | A | 20-04-2011 |
| 86 | F3AH3-T3 | Cattle pit #1 | At the pit's bottom | T3 | A | 20-04-2011 |
| 87 | F3BH1-T3 | Cattle pit #1 | Under the pit surface | T3 | B | 20-04-2011 |
| 88 | F3BH3-T3 | Cattle pit #1 | At the pit's bottom | T3 | B | 20-04-2011 |
| 89 | F3CH1-T3 | Cattle pit #1 | Under the pit surface | T3 | C | 20-04-2011 |
| 90 | F3CH3-T3 | Cattle pit #1 | At the pit's bottom | T3 | C | 20-04-2011 |
| 91 | F4AH1-T3 | Cattle pit #2 | Under the pit surface | T3 | A | 20-04-2011 |
| 92 | F4AH3-T3 | Cattle pit #2 | At the pit's bottom | T3 | A | 20-04-2011 |
| 93 | F4BH1-T3 | Cattle pit #2 | Under the pit surface | T3 | B | 20-04-2011 |
| 94 | F4BH3-T3 | Cattle pit #2 | At the pit's bottom | T3 | B | 20-04-2011 |
| 95 | F4CH1-T3 | Cattle pit #2 | Under the pit surface | T3 | C | 20-04-2011 |
| 96 | F4CH3-T3 | Cattle pit #2 | At the pit's bottom | T3 | C | 20-04-2011 |
| 97 | Fresh swine slurry | Swine slurry | - | - | - | 20-04-2011 |
| 98 | Fresh cattle's pit | Cattle's pit | - | - | - | 20-04-2011 |

### Cells fixation

Fecal aliquots have been diluted (1 :10w/v) in cold and filtered PBS buffer (130mM NaCl, 3mM NaH2PO4, 7mM Na2HPO4, pH 7.2; 4°C). In each tube, sterile glass pearls were added and homogenized by mechanical kneading for 3 min. and centrifuged at 700 rpm for 1 min. the aliquots have been diluted 1: 4 in paraformaldehyde-PBS (pH7.2). After 1 night incubation at 4°C, the aliquots have been centrifuged (8,000 rpm, 10 min.) and the aggregates of fixed cells have been washed with sterile PBS (1x, pH 7.2) then suspended in 50% ethanol-PBS (1x, pH 7.2) (1:1, v/v). The fixed cells are stocked in -20oC till further usage (Franks *et al*. 1998, Waldram *et al*. 2009). The cell fixation has been elaborated at the Environmental Health Laboratory, Dairy and Swine Research and Development Centre, Agriculture and Agri-Food Canada (Lennoxville, QC, Canada)

### The oligonucleotide probes

EUB338-I probe (Amann *et al*., 1990) was used to target the majority of the Eubacteria. The ARCH915 probe (Raskin *et al*., 1994) targets the Archaea. The control probe NON 338 (Waltner *et al*.,1993) was used as a negative control and has no target gene. Those probes are marked at the 5’ extremity with sulfoindocyanine Cy3 or Cy5. Six probes (Table 2) have been used for FISH detection. The marked probes were from Sigma Genosys (Sigma-Aldrich, Canada).

**Table 2.**
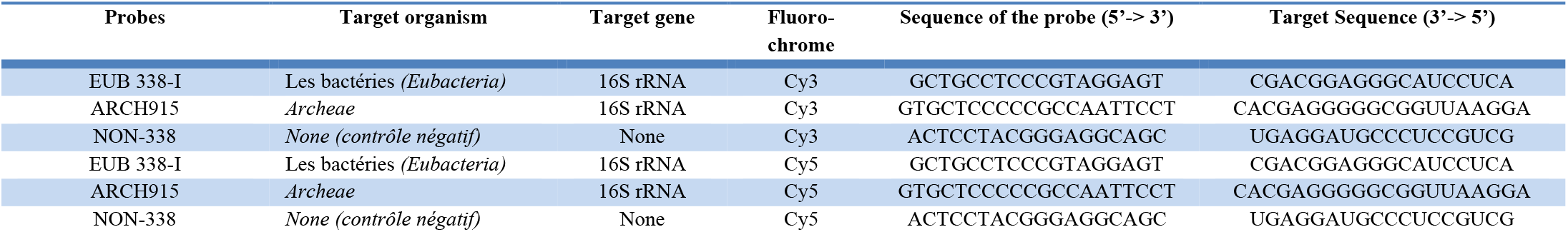
Sequences of rRNA-16S oligonucleotides showing the target regions of the used probes in the FISH technique to detect Bacteria and archeae.

### FISH

The fixed cells were diluted with 50% ethanol-PBS (1 :10 v/v) and spread on a surface of 1 cm^2^ on gelatin treated microscope slides. The cells were dried at room temperature for 30 min. to 1h. The slides were then dipped in 50%, 80% and 95% ethanol baths respectively for about 5 min each, this helps in the permeabilization of the cells as described by Lanthier *et al*. 2005; Takada *et al*. 2004 and Nakamura *et al*. 2009. 250 µl of acetylation solution (100 mM Triethanolamine, 0.25% acetic anhydride, 0.09% NaCl; pH 7.2) was added to each square (2 squares of 1cm^2^ per slide). The slides were incubated for 20 min at room temperature then rinsed with deionized water (Lanthier *et al*. 2002; Lanthier *et al*. 2005; Richter *et al*. 2007). The acetylation solution prevents the non-specific interaction between the fluorescent probes and the proteins. 100µl of hybridization buffer (35% deionized formamide, 0.9 M NaCl, 0.02 M Tris-HCl, 0.01% Sodium dodecyl sulphate; pH7.2) containing 0.25 ng/µl of each fluorescent probe were added directly to each square. The fluorescent probes were combined for each sample, 4 squares per 2 slides. To the first square, 100 µl of hybridization buffer containing 0.25 ng.µl^-1^ of EUB338-I probe marked with Cy3 and 0.25 ng.µl^-1^ of ARCH 915 probe marked with Cy5 have been added. To the second square, 100 µl of hybridization buffer containing 0.25 ng.µl^-1^ of EUB338-I probe marked with Cy5 and 0.25 ng.µl^-1^ of ARCH 915 probe marked with Cy3 have been added.

The negative control contained Cy3-labeled NON 338 and Cy5-labeled NON 338 probes at concentrations of 0.25 ng.µl-1 each. Autofluorescence was taken into account by adding 100µl of hybridization buffer without any fluorescent probe to the last square (Lanthier et al. 2002; Lanthier et al. 2005; Richter et al. 2007). The slides were then incubated in a hybridization dish moistened with paper towels saturated with 50 ml of hybridization buffer without fluorescent probes. Hybridization was performed, protected from light, using the “In slide out” oven (Boeckel Scientific, U.S.A) at 46°C for 2 h. After the hybridization step, the slides were washed at 48 °C in 2 baths of washing buffer (0.04M NaCl, 0.02M Tris-HCl, 0.01% sodium dodecyl sulfate, 0.005M EDTA; pH 7.2) and incubated for 20 min in the absence of light using the “Shake N’Bake” hybridization oven (Boeckel Scientific, U.S.A.) (Lanthier et al. 2002; Lanthier et al. 2005; Richter et al. 2007). The slides were rinsed with deionized water. A counterstain of 200 µl of DAPI (4’, 6’-diamino-2-phenylindole dihydrochloride; 10 µl.ml-1) was added to each square and incubated for 10 min at room temperature in the absence of light. After the slides were completely dry, coverslips were placed on each square after adding the fluorescence anti-bleaching agent (Prolong Gold, Invitrogen). The slides were allowed to dry overnight and were stored at room temperature and in the absence of light.

### Image acquisition and quantitative analysis

Slides were inspected under a Zeiss LSM 510 META confocal laser scanning microscope (Carl Zeiss, Germany) equipped with a Zeiss plan apochromat 63X oil immersion objective (numerical aperture of 1.4). For quantification, 20 microscopic fields taken randomly at different depths with the 63X objective were used. Images (resolution, 1,024 by 1,024 pixels, 8 bits pixels-1) were captured with Zeiss Axiocam camera (Zeiss, Germany). These images correspond to sections of 101.9 by 101.9 µm (area of 10,383.61 µm2; the total analyzed area is 207,672.2 µm2) and have a pixel size of 0.1 µm. The image acquisition program used is Zen 2009 Light Edition (Carl Zeiss Micro Imaging GmbH 2006-2009, Germany). Images were captured using the HeNe laser at 633nm for Cy5, DPSS laser at 561nm for Cy3 and Diode laser at 405nm for DAPI.

## Results and discussion

### Image Analysis

Double hybridization with Cy3-EUB338-I and Cy5-ARCH915 gave a better signal than with double hybridization Cy5-EUB338-I and Cy3-ARCH915 (Figure 1)

**Figure 1.**
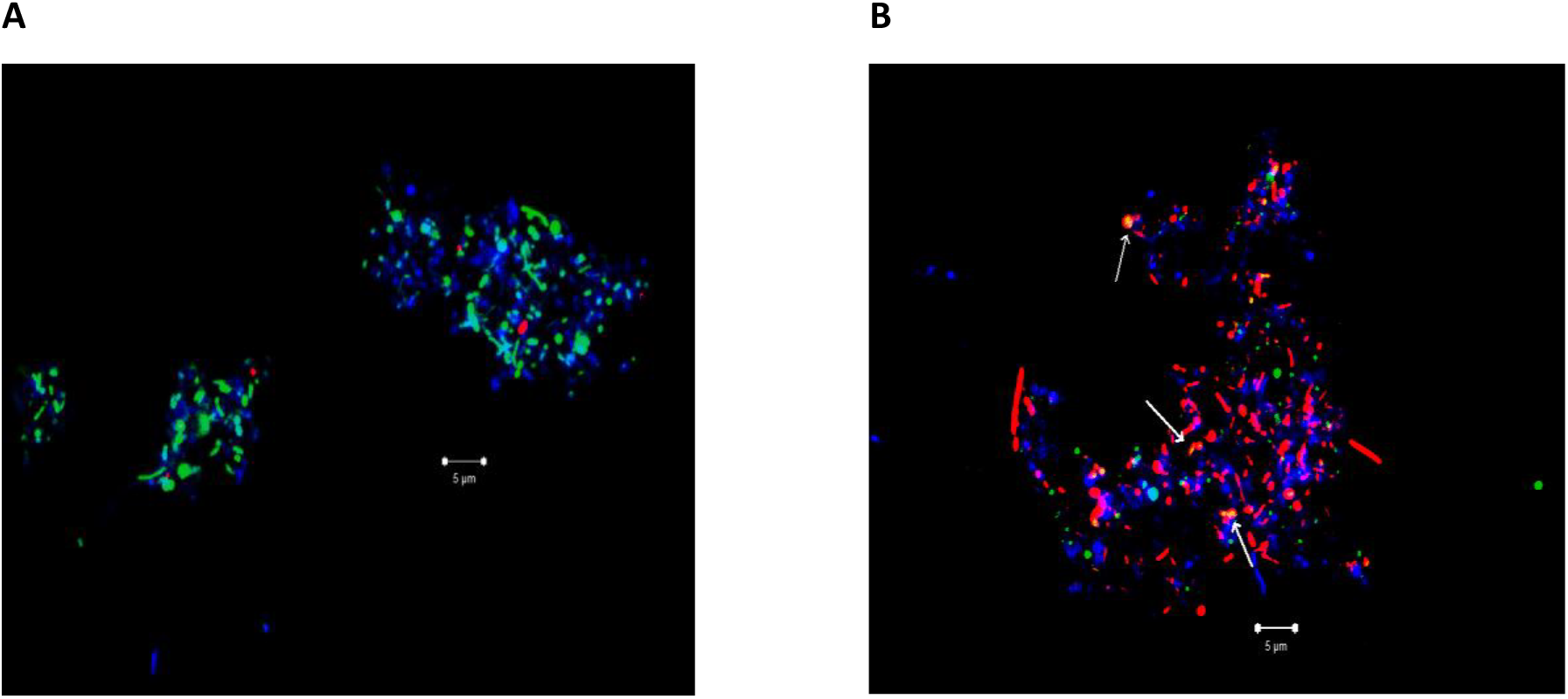
Confocal microscope laser scanning images of pig slurry #1 with double hybridization using the probes (A) Cy3-EUB338-I and Cy5-ARCH915 and (B) Cy5-EUB338-I and Cy3-ARCH915. Arrows indicate that cells were hybridized with both probes at the same time. Scale bar = 5 µm

The optimization of the FISH technique with the double hybridization Cy3-EUB338-I and Cy5-ARCH915 allowed to obtain good qualitative results in order to avoid the possibility of having colocalizations of the two probes in the cells. Thus, the significance of the quantification of bacteria and archaea in each of our samples would be greater. The quantification of bacteria and archaea present in the sample has been done by separating the 3 fluorescence images (DAPI, Cy3 and Cy5) with Zeiss 2009, then transforming them into binary images to quantify the pixels in each of the images by ImageJ, it is free software, which makes it difficult to find the appropriate plug-in.

### Potential and limitations of the FISH approach

We chose to use the standard FISH protocol using probes labeled at their 5’ end with either Cy3 or Cy5 and direct microscopic evaluation. An advantage of this approach is that the probes can be obtained commercially with a high degree of quality and relatively low cost. However, the high sensitivity of FISH probes did not allow the detection of all bacteria or archaea in our samples. It is assumed that the majority of microbial communities existing in the samples and having a low level of rRNA could not be detected (Oda et al. 2000).

## Conclusion

The objective of this study was to establish a double-staining FISH method to detect two types of microorganisms: bacteria and archaea. The images obtained show that there are more bacteria than archaea in the pig fecal sample studied. However, the choice of fluorescent probes for double-staining FISH is critical, as it is essential to use fluorochromes with a clearly differentiated fluorescent spectrum to create a unique spectral signature for each type of microorganism in the double-stained images. The fluorochromes Cy3, Cy5, and DAPI were chosen because they meet the aforementioned requirements and avoid signal colocalization.

## Dedication

In tribute to Dr. Martin Lanthier, supervisor of this work, who passed away (February 14, 2012), leaving the light of his works to shine within us, I dedicate this work to his soul.

## Acknowledgments

I would like to thank La cité collégiale for the academic support from all the teachers, and Dr. Guylaine Talbot and her team at the Dairy and Swine Research and Development Centre (Agriculture and Agri-Food Canada, Lennoxville) for their sampling efforts. I thank Dr. James Tambong (Eastern Cereals and Oilseeds Research Centre, Agriculture and Agri-Food Canada, Ottawa) for his supervision of this work, his support, encouragement, and invaluable advice. Special thanks to Michel Cloutier for his practical support, with great expertise, in the laboratory and to Denise Chabot for her assistance during image acquisition with the confocal laser scanning microscope.

